# Exploratory and prospective model of stent restenosis after Percutaneous Coronary Intervention using Multivariate Gaussian Subspatial Regression

**DOI:** 10.1101/445361

**Authors:** Victor Vicente-Palacios, Santiago Vicente-Tavera, P. Ignacio Dorado-Díaz, Antonio Sanchez-Puente, Jesús Sampedro-Gómez, Purificación Galindo, Itziar Gómez, José A. San Román, Francisco Fernández-Avilés, Pedro L. Sánchez

## Abstract

A new statistical analysis methodology, “Multivariate Gaussian Subspatial Regression” (MGSR), has been applied to randomized clinical trial data collected from percutaneous coronary intervention (PCI) patients, which combines the descriptive quality of Factorial Techniques and the predictive power of Gaussian Processes.

This model has been built from 3 different quantitative coronary angiographic core-lab measures of the same lesion from 2 separate angiograms (at baseline before PCI, at baseline immediately after PCI and at 12 months follow-up). Measurements of the pre-PCI variables of a patient are mapped to the factorial plane and predictions are visualized as regions of interest in this plane.

MGSR makes it possible to detect patients at risk of coronary stent restenosis or patients in whom ruling out the disease, in a graphical way; avoiding unnecessary, costly and possibly risky treatments for patients with no complications predicted; and advising to closely follow patients at risk. In addition, the model recovers missing values regardless of the variables, and once fitted, it corrects itself when more dependent variables are included.

MGSR software is freely available online at https://github.com/victorvicpal/MGSR.

## INTRODUCTION

Angioplasty and stenting are percutaneous coronary interventions (PCI) which have become routine practice for the revascularization of coronary vessels with significant obstructive atherosclerotic disease [1, 2]. Furthermore, PCI with stent implantation immediately performed is the gold-standard of treatment in patients with acute myocardial infarction, reducing the rates of death and recurrent ischemia as compared to medical treatment [3].

The long-term success of these procedures may be limited by restenosis [4, 5], a pathological process leading to recurrent arterial narrowing at the site of PCI which may manifest as a new myocardial infarction or force new-target-vessel revascularization. There is no equation attempting to predict which patients will develop restenosis, since there are probably multiple risk factors responsible for restenosis with anatomical vessel features and coexisting medical conditions. Thus, developing a model to explore and foresee the risk of restenosis would be a valuable tool for improving the stratification of patients, providing individually tailored follow-up and treatment.

Gaussian distributions are quite common in the medical and biological field. Angiography data are no exception and Gaussian processes are suitable to use. Gaussian processes use lazy learning and a measure of the similarity between points to predict new values accurately. Nevertheless, Gaussian processes rely on spatial or temporal domains, which are not present in this kind of data. We used a subspatial domain produced by dimensional reduction techniques to apply a Gaussian process such as cokriging. Thanks to this iteration we were able to forecast how PCI patients will behave after a PCI.

## MATERIALS AND METHODS

Stent Restenosis (SR) data were obtained from the previously published GRACIA-3 trial [6]. The GRACIA 3 trial is a randomized, open-label, multicenter, clinical trial that compares the efficacy of the paclitaxel-eluting stent with the conventional bare-metal stent. Patients with ST segment elevation acute myocardial infarction were enrolled from 20 Spanish hospitals.

To determine the incidence of SR, coronary angiography was performed at baseline and after 12 months of follow-up. All angiograms were analyzed at an independent angiography core laboratory (ICICOR, Valladolid, Spain) with a well-validated quantitative computer-based system (Medis, Leesburg, Va). The rate of SR was assessed by an experienced reader who was blinded to and not directly involved in the stent-implantation project.

Angiographic follow-up at 12 months was completed in 299 out of 346 (86%) randomized and eligible patients. SR was defined following the rate of binary restenosis after 12 months of angiographic follow-up. Binary angiographic restenosis was defined as a >50% narrowing of the lumen diameter in the target segment (defined as all portions of the vessel that received treatment within the stent zone, including the proximal and distal 5-mm margins) [7].

The method chosen to perform the analysis is the Multivariate Gaussian Subspatial Regression (MGSR) [8] model(1). This method combines factorial techniques and Gaussian processes. Factorial techniques gives us a description of our data set in terms of unobserved variables called factors, that in this case will be interpreted as coordinates in a 2D subspace. On the other hand, Gaussian processes are random structures that possess known statistical properties. As we are dealing with subspatial structures and our intention is to visually explain our data, cokriging methods also known as Gaussian process multivariate regression will be used. The Cokriging method predicts the value of a point in space given the similarity between surrounding points; which in our case will allow us to complete the picture in the regions where we have no data points and also to recover the original variables for any point given its subspatial coordinates.

A new patient, for which we only have the pre-PCI variables measured, can also be projected into this subspace using the factorial techniques. And then recover the post-PCI variables using Gaussian processes and the similarity with the rest of the data points. Although this is the natural case (projecting the pre-PCI variables and predict the post-PCI ones), this method let us predict any missing variables from the ones we have.

This method had the advantage of combining both descriptive and predictive techniques, which makes it very flexible. We can use the predictive power of this model to delimit relevant regions in our descriptive 2D subspace: any new patient would correspond to a point in this subspace and be automatically categorized in terms of its coordinates, and then we can obtain the prediction for the post-PCI variables. Afterwards, we can follow the evolution of each patient as the projection changes when we add the post-PCI variables (ideally, if the original predictions match exactly with the measurement of the post-PCI variables, there would be no shift in the projection).

### Multivariate Gaussian Subspatial Regression

We first briefly describe the MGSR[8] model(Fig. 1) and summarize its statistical properties.

Let ***X_N ×P_*** be the data matrix that is composed of *P* variables and *N* individuals. We chose a Classical Biplot [9] as a dimensional reduction technique. Biplot is a graphical representation of the data matrix ***X*** as ***X*** ≈ ***RC^T^***, where ***R*** and ***C*** are *N* × 2 and 2 × *P* matrices known as the row and column coordinates respectively. From now on we will refer to ***R*** as the subspatial coordinates. Each individual can be represented as a point in a scatter plot in those coordinates. These representations allows researchers to simply visualize and understand their sample data. Nevertheless if we need to look beyond and build a predictive model, biplots are limited.

To overcome this issue we will apply a Gaussian Process over the resulting Biplot plane. We might consider our first factorial plane as if it were a mining region in which points would represent our exploratory drilling sample. Multivariate geostatistical procedures [10] would help us to forecast where it would be better to drill in the unknown region depending on the desired mineral we want to extract. In our case we do not aim to “drill” but to recover the variables of a hypothetical individual which lies in an unknown region, that is, a blank space of the biplot plane not covered by our sample data. On this basis, we propose a cokriging exploration of the biplot plane.

Previous to applying cokriging we need to compose a suitable matrix that will be used later on. For this purpose, we proceed to standardize by columns the ***X**_N ×P_* by removing the column means and dividing by its standard deviation. Then, we associate the obtained coordinates ***R*** with the standardized matrix obtaining ***Z***(*u*) = [***X*** (***R***)] where *u* = < ***r_s_*** >,*u* ∈ *S* are the set of subspatial locations and *S* represents the generated supspace.

**Fig 1.**
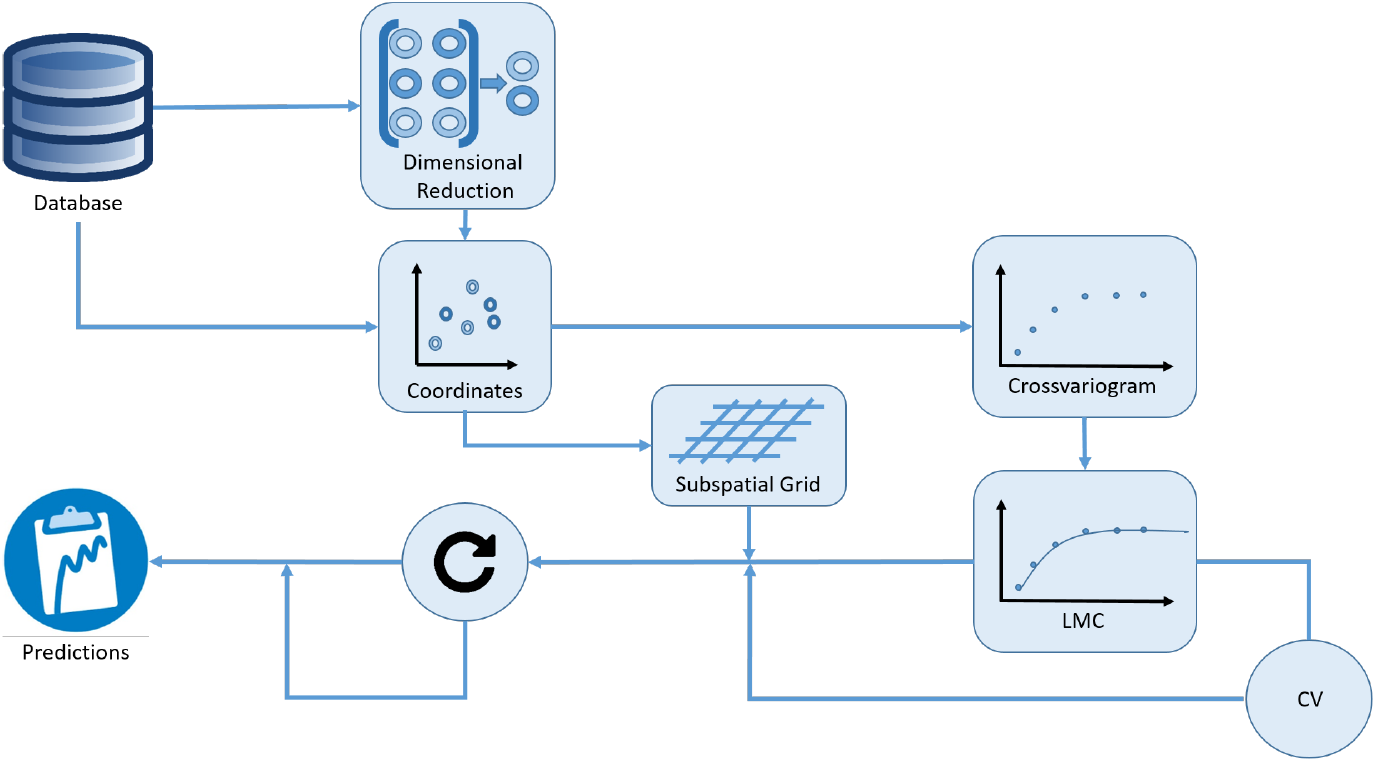
MGSR algorithm

Subsequently, we build an experimental subspatial cross-variogram [11] Γ<_*i,j*_(*h*), where *h* is the distance between points and *i, j* represent our features, that encode the correlation between variables at two different points given the distance between them.

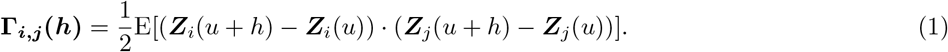

Using a more feasible approach, we calculate each experimental cross-variogram 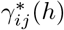 as

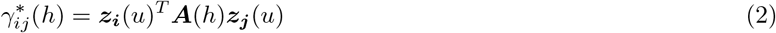

where ***z_i_*** = (***Z_i_***(*u*_1_),…, ***Z_i_***(*u_n_*))^*T*^ is the data vector for the random field ***Z_i_*** and ***A***(*h*) is the spatial design matrix. Hence, if *i* = *j* we obtain a direct cross-variogram.

The spatial design matrix ***A***(*h*) is calculated as

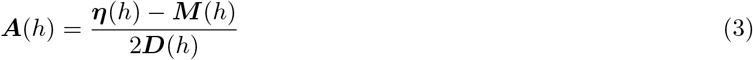

where *M*(*h*) is the *N* × *N* binary matrix *m_uv_* with *m_uv_* = 1 if the vector linking sampling locations *u* and *v* corresponds to lag *h* and *m_uv_* = 0, otherwise; ***η***(*h*) is the *N × N* diagonal matrix *η_uv_* with 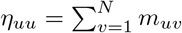 (i.e., the number of neighbors for sampling location u at lag h), and ***D***(*h*) is the number of pairs of observations at lag *h*.

Therefore, we obtain *P*(*P* + 1)/2 experimental cross-variograms that need to be fitted. Before introducing the procedure to fit the experimental cross-variograms, it is necessary to understand the different elements which make up the whole:

- Nugget: represents variability at small distances (*h* ≈ 0).
- Sill: The semivariance value *b* at which the semivariogram levels off.
- Range: The *a* distance at which the semivariogram reaches the sill value.

We have chosen the Linear Model of Coregionalization (LMC) to fit our set of cross-variograms. The LMC can be expressed as a multivariate nested semivariogram model [10].

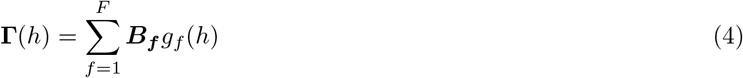

where **Γ**(*h*) is the *P × P* matrix of semivariogram values at lag *h*, and ***B_f_*** is the *P × P* matrix of sills [12] of the basic semivariogram function *g_f_*(*h*). ***B_f_*** has to be positive semidefinite [12, 13] to ensure that the variance-covariance matrix is also positive semidefinite. Although different approaches of LMC can be found in the literature, we use the algorithm defined by Pelletier et al. [14] owing to its clarity:

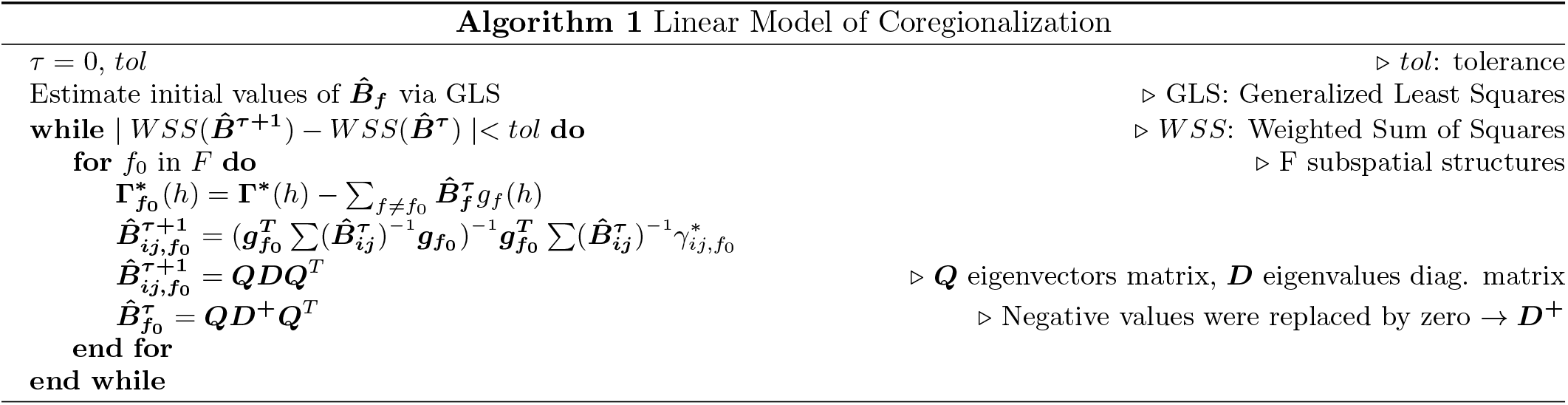

Unlike geostatistical analyses [15], there is no real field where boundaries restrict our study. However, this aspect is more positive than negative because we are able to create a simpler grid without losing information.

By establishing the interval between the maximum and minimum location in their different dimensions we create a frame, which can be extended if necessary. Subsequently, we build the grid choosing a suitable division.

Using simple cokriging [10] we are able to project our predictions onto the grid, and thus are able to compare the results within the variables.

Cokriging is the multivariate extension of kriging, whose main purpose is to compute a weighted average of the sample values in close proximity to the grid point. It searches for the best linear unbiased estimator, based on assumptions on covariances.

Simple cokriging is based on three assumptions: stationary, known means and known covariance functions. The simple cokriging estimator is

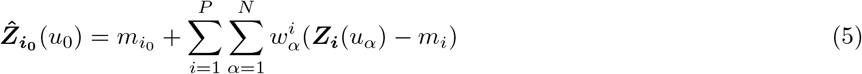

where *u*_0_ is the grid location and *u_α_* the sample location, 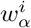 is the weight and *m* corresponds to the means of our variables. To estimate the 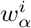 we use ***C_uv_****w_i_* = ***c_ui0_*** [16] as estimators, where ***c_ui0_*** is the *N × N*_0_ covariance matrix between grid and sample locations and ***C_uv_*** is the *N × N* covariance matrix; being *N*_0_ the total number of points in our grid. We can also describe this relation as

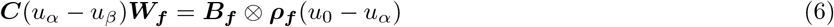

where *ρ_f_*(*h*) represents the covariance function and *u_β_* a sample location. ***C_uv_****w_i_* can also be expressed as

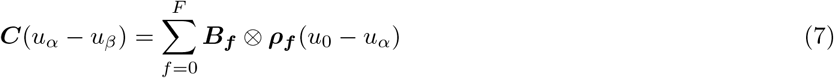

It is worth noting that both covariance matrices were determined in a way similar to the theoretical semivariogram; however, we cannot assume that both approaches were identical [10].

Once we obtained ***W***; as we were using a standardized matrix, all means were zero, and the estimation of 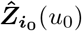 was direct

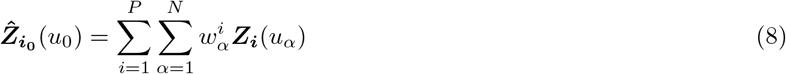

The cokriging error is determined as

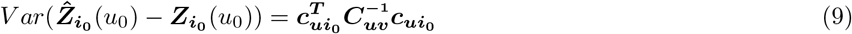

As soon as we obtained the matrix 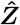 we were able to make predictions over the grid.

Let ***Y*** be the input or known *N*\* × (*P* – *P*\*) matrix and ***Y***\* the *N*\* × *P*\* matrix, where *P*\* are the variables to predict 1 < *P*\* < *P* and *N*\* the number of elements to predict. If ***m*** is the distance vector between 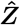 and ***Y*** such as 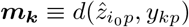 where *p* =1,…, (*P* – *P*\*) and *k* =1,…, *N*\* then ***m*** provided the positions on the grid where we could uncover the 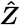 predictions.

An alternative explanation of the model can be found at [8].

## RESULTS

Our aim was to develop an exploratory and prospective model of the obstructed diameter of the artery for two different conditions, post and follow-up PCI in a simpler, visual and intuitive way. For this matter we had 3 pre-PCI variables (Legth, obstructed diameter and reference diameter of the artery) and 2 post-PCI variables (immediately-post and 12-month follow-up PCI diameters).

Our starting matrix was composed of 337 patients and 5 variables; however, there were only 180 patients without missing values (Table I), representing 53% of the total.

**TABLE I.** Stent data Summary (mm.)

| Variables | Pre |  |  | Post | Follow-Up |
| --- | --- | --- | --- | --- | --- |
|  | Seg Length | Obs Diam | Ref Diam | Obs Diam | Obs Diam |
| mean | 43.42 | 0.78 | 2.89 | 2.35 | 2.22 |
| S.E. | 1.16 | 0.03 | 0.04 | 0.04 | 0.04 |
| max | 111.98 | 1.91 | 4.32 | 3.68 | 3.53 |
| min | 8.86 | 0.00 | 1.55 | 1.09 | 0.46 |
| Fig. ref. | Seg.Length.PRE | Obs.Diam.PRE | Ref.Diam.PRE | Obs.Diam.POST | Obs.Diam.FU |

Classical Biplot [9] is the initial dimensional reduction procedure in our method, in particular we chose Row Metric Preserving Biplot. Biplots represent matrix elements as points and variables as vectors (Fig. 2). The inertia absorption of the first two factor axes was 69%. Despite of the loss of information, we kept two first axes in order to facilitate the visualization and simplify the cokriging iteration later on.

As seen in Fig.2 points represent our patient’s cohort and vectors describe the behavior of the analyzed variables. Once we got our Biplot coordinates, we calculated the experimental cross-variogram which describes the degree of subspatial dependence of the calculated factorial plane. Experimental cross-variograms are represented in Fig. 3 as a set of scatter-plots. The distribution of the experimental cross-variograms follows a power distribution *g_f_* (*h*) = *h^a^*, which has been the chosen function for the Linear Model of Coregionalization (LMC) fitting (*a* = 1.83). Unlike continuous fields, subspatial domains do not have verges. As a result of this lack of boundaries and the linear response of classical Biplot, Power or Linear distributions are more appropriate than Gaussian or Spherical ones. We chose a Power distribution instead of a Linear one due to a slightly better fit.

Table II displays the results of the LMC.

In addition, we calculated the root mean square error (RMSE) of our model [10]. The cross-validation method was based on a common resampling. Each sample value ***Z***(*u_α_*) was removed in turn from the data set and a value 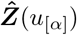 was estimated at that location using the *N* – 1 other samples. As a result we determined the residuals 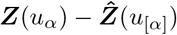 of our model.

**Fig 2.**
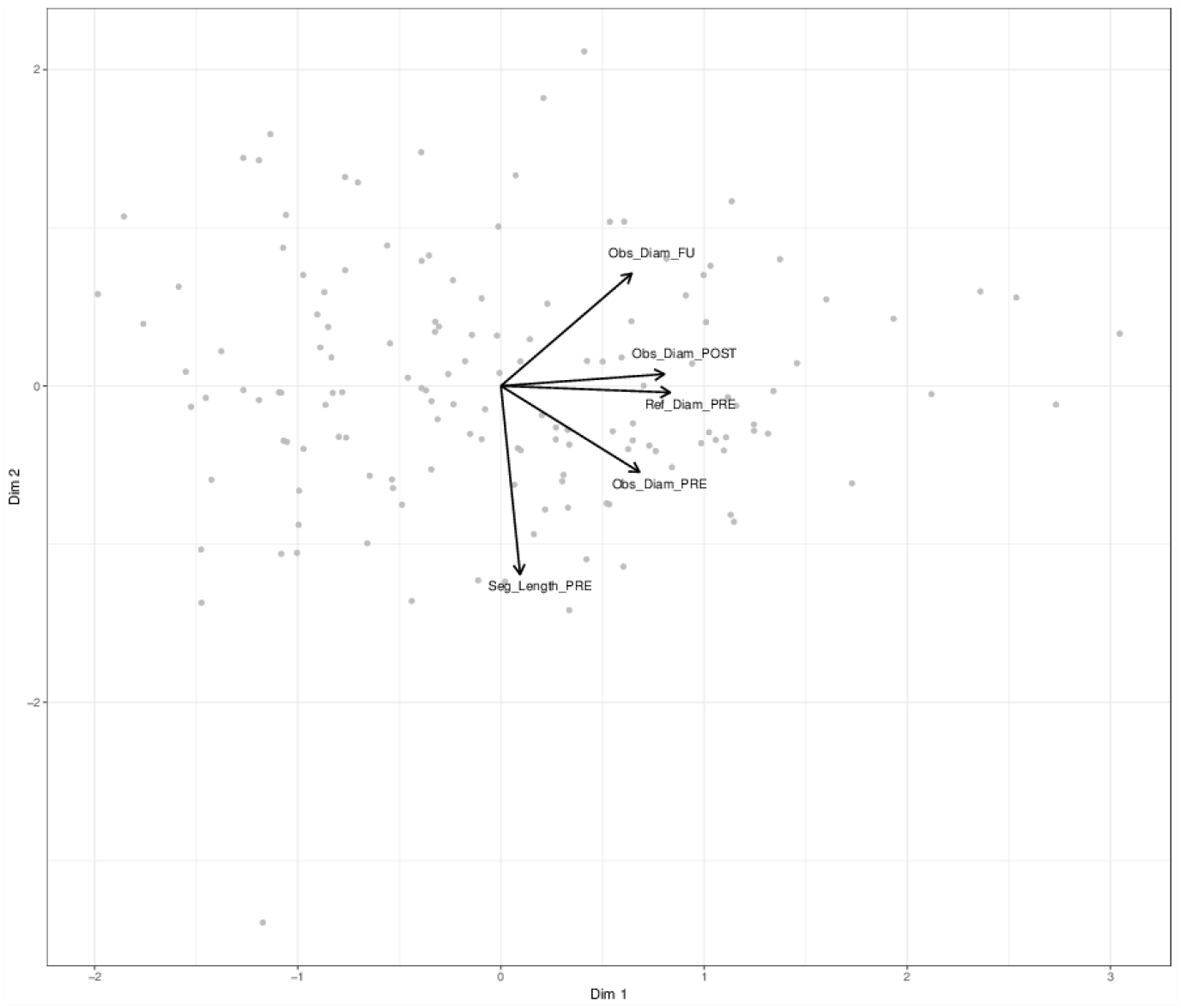
JK-Biplot

**Fig 3.**
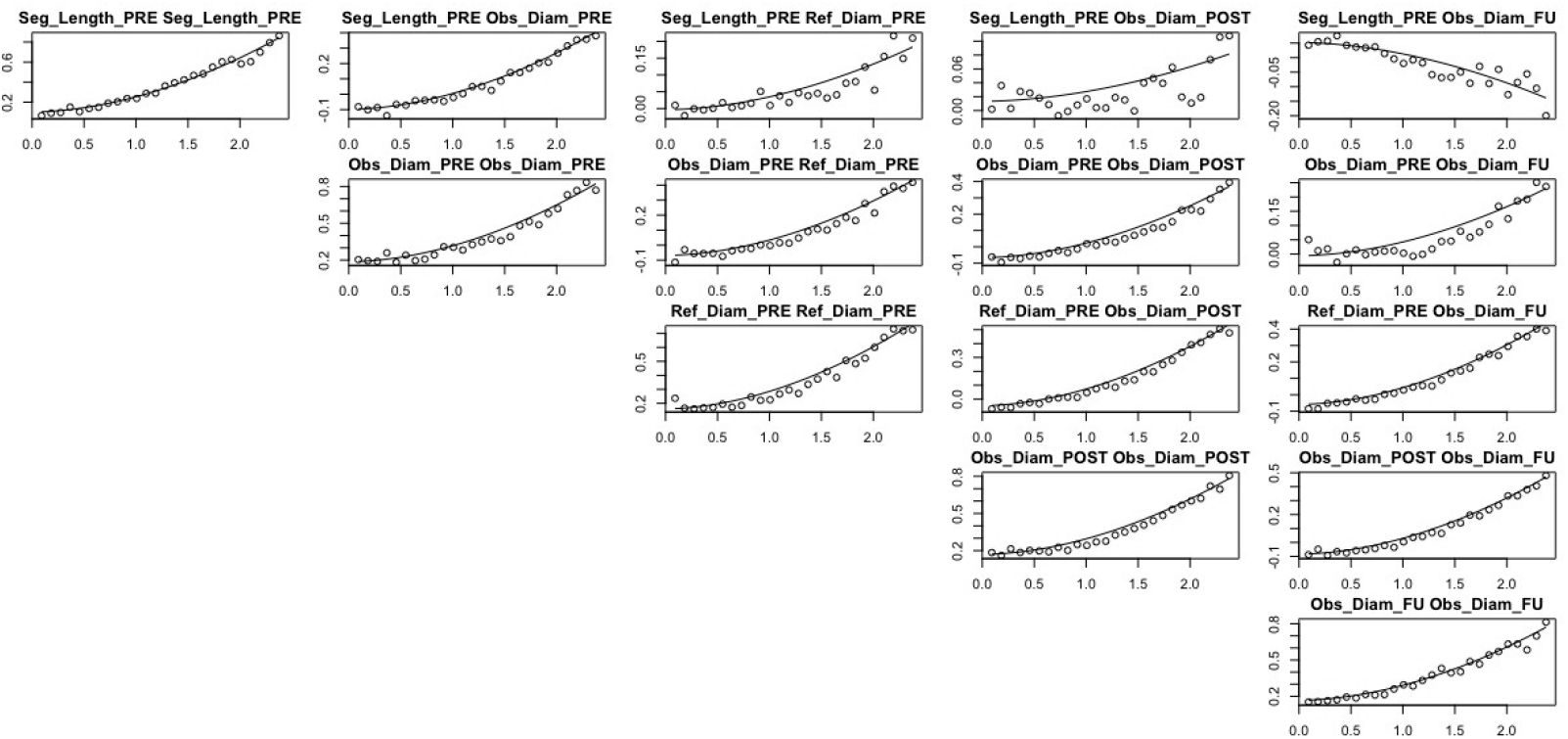
Cross-Variogram

Also, we compared our results with classical Multivariate Regression (MVR) models by applying the same kind of cross-validation. To be able to compare MGSR and MVR we needed to fit 10 different multivariate models. All of these models had 3 input variables and 2 output variables in such a manner that each variable was considered as a dependent variable 6 times and as independent, 4 times. Table III shows the RMSE of MGSR and the 10 MVR models. Empty values on the table III correspond to input variables.

**TABLE II.**
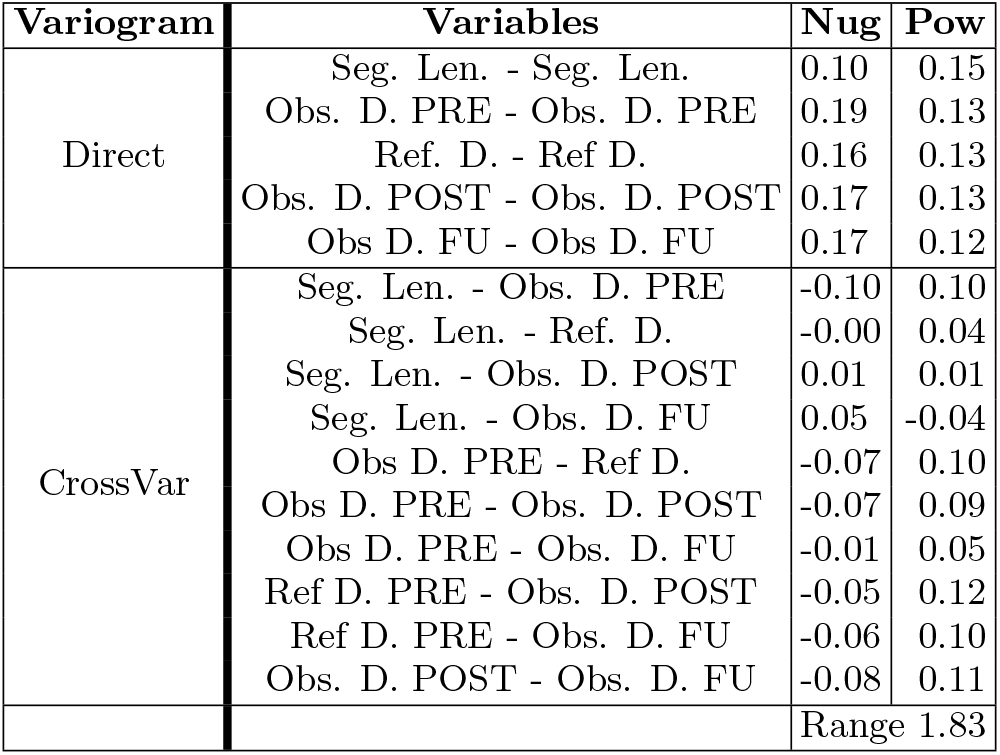
Summary of LMC results

| Variogram | Variables | Nug | Pow |
| --- | --- | --- | --- |
| Direct | Seg. Len. - Seg. Len. | 0.10 | 0.15 |
|  | Obs. D. PRE - Obs. D. PRE | 0.19 | 0.13 |
|  | Ref. D. - Ref D. | 0.16 | 0.13 |
|  | Obs. D. POST - Obs. D. POST | 0.17 | 0.13 |
|  | Obs D. FU - Obs D. FU | 0.17 | 0.12 |
| CrossVar | Seg. Len. - Obs. D. PRE | -0.10 | 0.10 |
|  | Seg. Len. - Ref. D. | -0.00 | 0.04 |
|  | Seg. Len. - Obs. D. POST | 0.01 | 0.01 |
|  | Seg. Len. - Obs. D. FU | 0.05 | -0.04 |
|  | Obs D. PRE - Ref D. | -0.07 | 0.10 |
|  | Obs D. PRE - Obs. D. POST | -0.07 | 0.09 |
|  | Obs D. PRE - Obs. D. FU | -0.01 | 0.05 |
|  | Ref D. PRE - Obs. D. POST | -0.05 | 0.12 |
|  | Ref D. PRE - Obs. D. FU | -0.06 | 0.10 |
|  | Obs. D. POST - Obs. D. FU | -0.08 | 0.11 |
|  |  | Range | 1.83 |

**TABLE III.** RMSE MGSR and Multivariate Regression

| Algorithm | PRE |  |  | POST | FU |
| --- | --- | --- | --- | --- | --- |
|  | Seg. Length | Obs. Diam. | Ref. Diam. | Obs. Diam. | Obs. Diam. |
| **MGSr** | 0.19 | 0.40 | 0.33 | 0.35 | 0.32 |
| **MVR1** | - | - | - | 0.43 | 0.50 |
| **MVR2** | - | 0.38 | - | 0.43 | - |
| **MVR3** | - | 0.38 | - | - | 0.49 |
| **MVR4** | - | 0.38 | 0.43 | - | - |
| **MVR5** | - | - | 0.46 | 0.46 | - |
| **MVR6** | - | - | 0.44 | - | 0.50 |
| **MVR7** | 14.47 | - | 0.42 | - | - |
| **MVR8** | 14.97 | 0.39 | - | - | - |
| **MVR9** | 14.40 | - | - | 0.42 | - |
| **MVR10** | 14.89 | - | - | - | 0.50 |

### Predictive value of the model

At this point we used cokriging to make predictions over the blank regions of fig. 2, where we do not have data points corresponding to real patients. We built a rectangular grid containing all the Biplot projections. Grey points in figs. 4 and 5 show the grid distribution where we applied a Cokriging iteration. Owing to computational issues, we chose a 0.1 distance between grid points. A tighter choice would not improve results.

We illustrate our model as if it were a map of elements (Fig. 4). Each point represents an hypothetical patient with different characteristics. Patients nearby tend to be more similar than those that are farther apart. We can use the predictions from the cokriging iteration to delimit regions of interest in this plane. Our main interest is the stent restenosis region (in red), that we have defined as the region in which the predicted lumen diameter at follow-up is less than 50% of the reference diameter, in which patients should be watched closely. We have also highlighted the regions of large acute gain (in green) and low acute gain (in blue), that have been defined as the regions in which the predicted lumen diameter right after the intervention is greater than 83% of the reference diameter and less than 75% of the reference diameter, respectively.

Cokriging errors were also calculated and represented in Fig. 5. The regions in salmon color have the most accuracy, meanwhile the regions in white and the regions ranging from yellow to green are less accurate. We can see that the predicted points near the origin of the coordinate system are less accurate than the surrounding points [9], this effect corroborates the Biplot quality of rows representation theory [17]. It is worth noting that the regions of greater interest in this study (restenosis, low acute gain and large acute gain) lie in the region where the model is most accurate.

To illustrate how MGSR can be used, we present a couple of examples over fig. 4. In patient number 1, pre-PCI MGRS predicts that the development of stent restenosis (red area) is uncommon to happen. This is confirmed in the post-PCI MGRS model predicting an optimal evolution, which is corroborated at the 12 months angiographic follow-up. In patient number 1, we could avoid repeated clinical visits to the doctor as optimal evolution is predicted.

**Fig 4.**
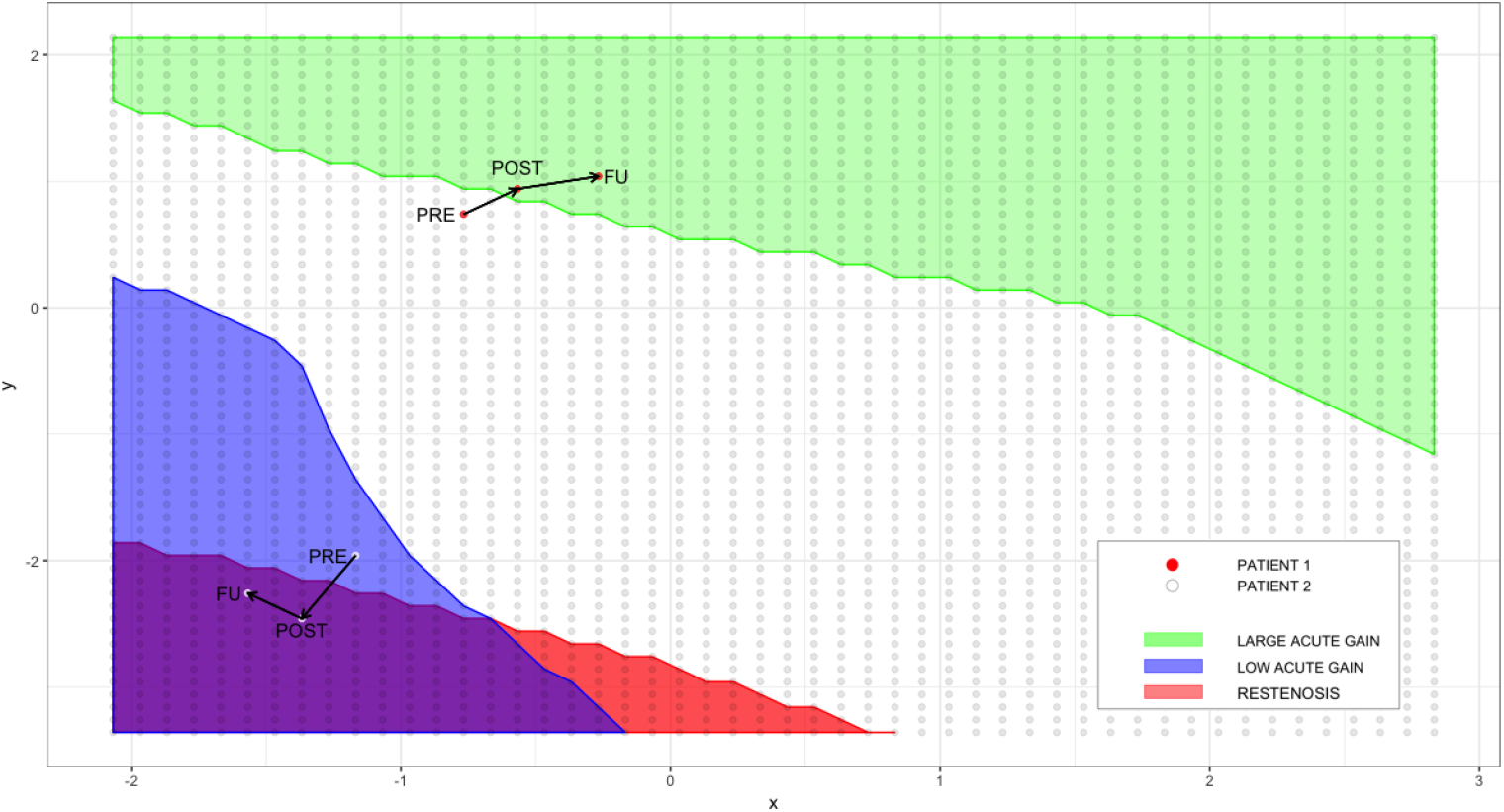
MGSR model

**Fig 5.**
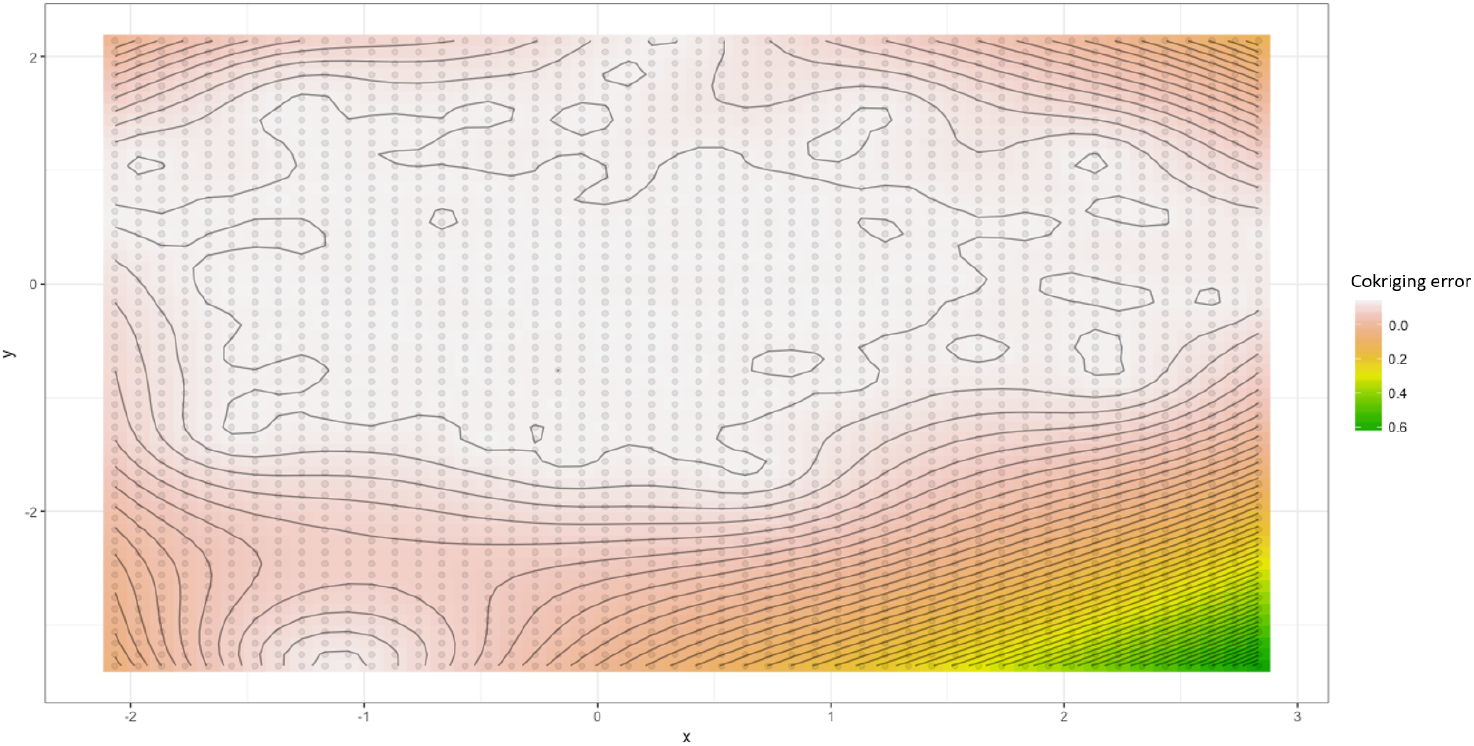
Cokriging error

Furthermore, we could also avoid double antithrombotic therapy continuation after 12 months, as no stent complications are predicted.

In contrast, patient number 2 pre-PCI prediction is close to the red area of stent restenosis. The risk of stent restenosis MGRS prediction is even worse immediately post-PCI and stent restenosis is confirmed at the 12 months angiographic follow-up. In patient number 2, we should improve the post-PCI result from the predicted one: we should recommend a close clinical follow-up, a 12 month coronary angiography should be performed, and double antiaggregation should not be discontinued as we predict future adverse events.

These estimations can be done not only in a chronological way, as seen in Fig. 4 but also to calculate missing values. For instance, if the reference diameter of the artery was not measured on a patient, we could estimate its value without fitting a new model by searching over our model (Fig. 4) the most similar hypothetical patient based on the real patient’s known values.

Almost half of our dataset contains missing values which can be calculated as we explained above. A summary of our outcomes are shown in Table V.

**TABLE IV.**
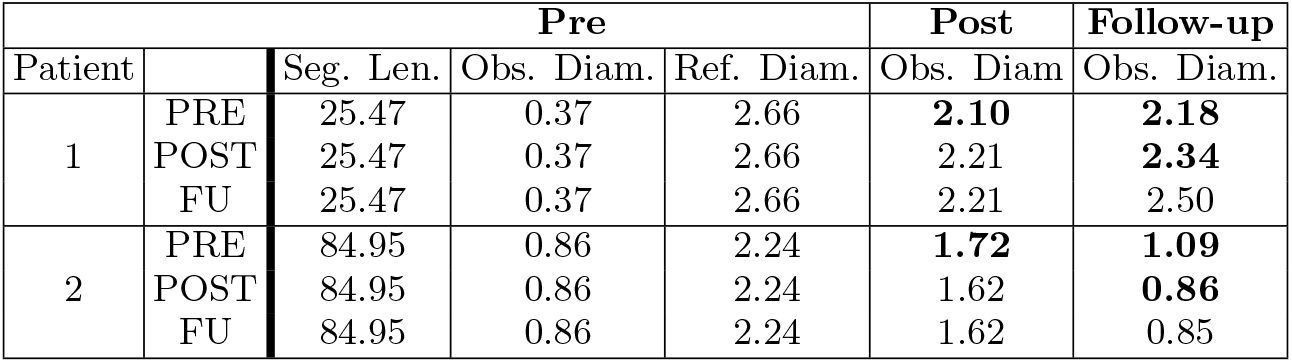
Examples of prediction

**TABLE V.**
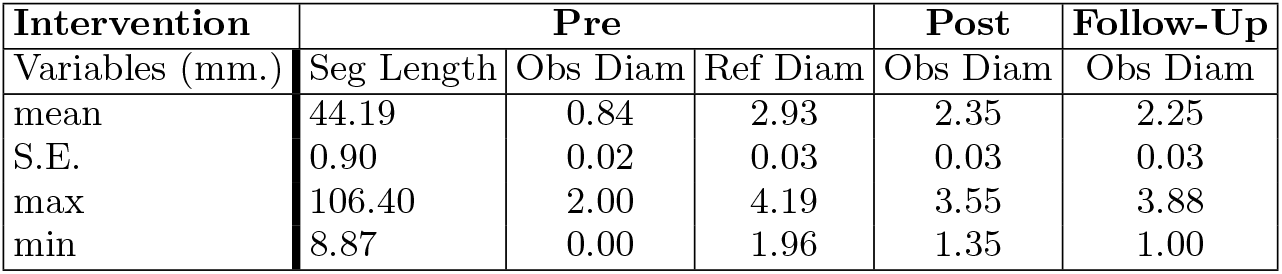
Missing Values

## DISCUSSION

### Methodological discussion

The proposed model is very versatile and flexible for studying continuous variables from an exploratory and prospective point of view. These variables can be combined according to the needs of the researcher. We have also demonstrated that by combining the knowledge of the domain expert and the visualization power of our method, we are able to create straightforward visualizations.

Besides its versatility, we were able to forecast the evolution of a specific variable in different time moments. In our case, the analyzed variable was the obstructed diameter (PRE, POST and Follow-up PCI) which is the most important feature to quantify possible future patient outcomes. This characteristic was already reported in papers previously published by some of the authors of this work[8].

We proved that by combining two different procedures we could obtain interesting outcomes. Factorial techniques benefit from Gaussian Processes without losing their exploratory attributes and Gaussian Processes can be applied over a non-continuous field. We can also reproduce our model in a visual way letting researchers better understand their data. We strongly believe that our model is extensible to other bioscience fields such as pharmacy, biology or ecology among others.

In addition, missing values, no matter which variable, can be estimated. This makes it possible to estimate lost information from the trial, where missing values are habitual. This fact increases the chances of having selection bias while analyzing the data. By allowing researchers to partially recover this missed information we could decrease it. Nonetheless, this potential benefit relies on high goodness of fit of the model and on a sufficiently representative sample.

Although the strengths are promising, some limitations might be pointed out. Currently, MGSR is limited to continuous data making difficult the analysis of ordinal or categorical variables among others. Also, as seen in the results section by only keeping two dimensions in the dimensional reduction step we risk the loss of information that could penalize the next steps. To overcome this issue we should consider extending our analysis to multiple dimensions. In our research, we had only used linear Factorial techniques. Further investigation is required into the opportunities for using non-linear procedures.

Besides these limitations, MGSR needs to be improved in several practical aspects. For instance, the computation time needed can be too large with a large set of variables. This is an inherited problem of multivariate spatial Gaussian Processes. Also, to fit an MGSR model we need to follow a series of steps that can be tedious for users. In order to improve both issues, we intend to better automate the LMC fitting to be more user-friendly and try different matrix factorizations or Bayesian approaches to reduce time on the Cokriging iteration.

In terms of the application, another limitation should be mentioned. The size of our data was not big enough to split them in a train and test datasets. We could have better proved the robustness of our model if we had had a larger sample.

In conclusion, MGSR has a particular methodological opportunity which is definitely worth investigating.

### Clinical discussion

Currently, PCI is the most common procedure to treat coronary atherosclerotic obstructions. The total number of PCI performed in Spain in 2015, with approximately 45 million inhabitants, were 67,671 procedures, with a ratio of 1,466 per million inhabitants. In addition, 18,418 were carried out during the acute phase of myocardial infarction (21.7%) [18].

The Achilles tendon of PCI is restenosis, which leads to recurrent arterial narrowing at the site of intervention. Restenosis is caused by chronic inflammatory disease featuring the activation of immune cells and excessive growth of vascular smooth muscle cells within the injured vessel wall. When PCI was initially carried out, in the late 1970s, angioplasty was limited to isolated balloon inflations. However, and although the artery would initially be opened successfully using a balloon, approximately 30% of all coronary arteries began to close up some months after balloon angioplasty owing to restenosis [19]. By the mid-1980s professionals began to design new devices to improve the durability of PCI procedures, avoiding restenosis. One such device was the stent, a metal tube that is inserted after balloon inflation, thus preventing negative remodeling of the treated vessel, the major cause of restenosis after conventional balloon angioplasty. But, while metal stents eliminated many abrupt artery closures, restenosis persisted in about 20% of cases [20]. In the early part of this century, the solution to restenosis moved away from mechanical devices towards the use of pharmaceuticals. Professionals started to test a variety of drugs that were known to interrupt the biological processes that cause restenosis. Stents were coated with immunosuppressive and anti-proliferative drugs, clinical trials began, and the era of the drug-eluting stents commenced. We are now in the second generation of drug eluting stents and restenosis has been reduced to 10%. However, and considering the increasing numbers of PCI performed worldwide and the widespread use of drug eluting stents, restenosis is still a health care problem of magnitude and identification of patients at risk is an important undertaking [21].

In this regard, the model to predict the risk of restenosis for improved stratification of patients proposed in our study is of clinical and economic relevance. Following this model, we could prescribe bare metal stents and decrease costs for patients at low-moderate risk of restenosis and improve outcomes in patients at high-risk of restenosis by implanting drug-eluting stents and performing established 12 month follow-up catheterization to rule out restenosis and avoid adverse clinical outcomes. Although restenosis may be associated with a recurrence of stable angina symptoms, it is recognized that up to 1 out of 3 patients present with myocardial infarction or unstable angina amenable to repeat catheter intervention [22].

The GRACIA-3 trial is unique in developing this predictive model for several reasons. First of all, it provides the opportunity to compare bare metal stents versus drug eluting stents in a clinical context, ST elevation acute myocardial infarction, where the benefit of new drug eluting stents is still discussed and under comparison [23]. Thus, our predictive model would be specifically useful in this specific scenario. Second, it is important to put into perspective the number of paired angiographic studies performed in the GRACIA-3 trial (70%), which surpasses surveillance studies in large cohorts of patients [24]. Thus, it will be difficult to find a cohort of patients so well characterized to develop and test a predictive model. Finally, the data come from a randomized clinical trial where all angiograms were analyzed at an independent angiographic core laboratory (ICICOR, Valladolid, Spain) with a gold-standard validated quantitative computer-based system (Medis, Leesburg, Va). Angiographic restenosis defined as binary angiographic restenosis > 50% narrowing of the lumen diameter in the target segment was assessed by an experienced reader who was blinded to treatment assignment and clinical outcome. Thus, the methodology used to define the presence of restenosis was of maximum rigor and quality.

## Notes

#### Summary of Updates

Sections "Materials and methods"; "Results" and "Discussion" updated;

